# Lipid-derived electrophiles induce covalent modification and aggregation of Cu,Zn-superoxide dismutase in a hydrophobicity-dependent manner

**DOI:** 10.1101/740688

**Authors:** Lucas S. Dantas, Lucas G. Viviani, Alex Inague, Erika Piccirillo, Leandro de Rezende, Graziella E. Ronsein, Ohara Augusto, Marisa H. G. de Medeiros, Antonia T.-do Amaral, Sayuri Miyamoto

**Author notes:** Corresponding author: Sayuri Miyamoto, Departamento de Bioquímica, Instituto de Química, Universidade de São Paulo, Avenida Professor Lineu Prestes, 748, Bloco 10 Superior, sala 1074, São Paulo, SP, Brazil 05508-000. Abbreviations HHE - 4-hydroxy-2-hexenal; HNE - 4-hydroxy-2-nonenal; HEX - Hexenal (*trans*-2- hexen-1-al); NON - Nonadienal (*trans,trans*-2,4-nonadienal); DEC - Decadienal (*trans,trans*-2,4-decadienal); Seco-A - 3β-hydroxy-5-oxo-5,6-secocholestan-6-al; Seco- B - 3β-hydroxy-5β-hydroxyB-norcholestane-6β-carboxyaldehyde; SOD1 - cooper, zinc-superoxide dismutase; ALS - amyotrophic lateral sclerosis; SEC - size exclusion chromatography; LC-MS/MS - Liquid Chromatography Coupled to Tandem Mass Spectrometry.

## Abstract

Lipid peroxidation generates a huge number of reactive electrophilic aldehyde products. These reactive aldehydes can modify macromolecules such as proteins, resulting in loss of function and/or aggregation. The accumulation of Cu,Zn-superoxide dismutase (SOD1) aggregates is associated with familial cases of amyotrophic lateral sclerosis (ALS). Recent studies have shown that lipid and its oxidized derivatives may play a role in this process. Here we aimed to compare and characterize the ability of lipid-derived electrophiles with different hydrophobicities to induce SOD1 modification and aggregation *in vitro*. SOD1 was incubated with 4-hydroxy-2-hexenal (HHE), 4-hydroxy- 2-nonenal (HNE), 2-hexen-1-al (HEX), 2,4-nonadienal (NON), 2,4-decadienal (DEC) or secosterol aldehydes (Seco-A or Seco-B) at 37°C for 24 h. Size exclusion chromatography analysis showed that hydrophobic aldehydes smarkedly enhances apo- SOD1 aggregation. More importantly, aggregation level was positively correlated to calculated aldehyde hydrophobicities (LogP). Protein sequencing by LC-MS/MS showed that aldehydes covalently modifies SOD1 at aggregation prone regions. For instance, specific lysine residues located mainly nearby the dimer interface (K3, K9) and at the electrostatic loop (K122, K128, K136) were ubiquitously modified by all aldehydes. The α,β-unsaturated aldehydes also promoted modifications on histidine and cysteine residues, with H120 and C6 being the most commonly modified residues. Overall, our data suggest that electrophile’s hydrophobicity is a critical factor that strongly influences protein aggregation propensity.

**Graphical abstract:** 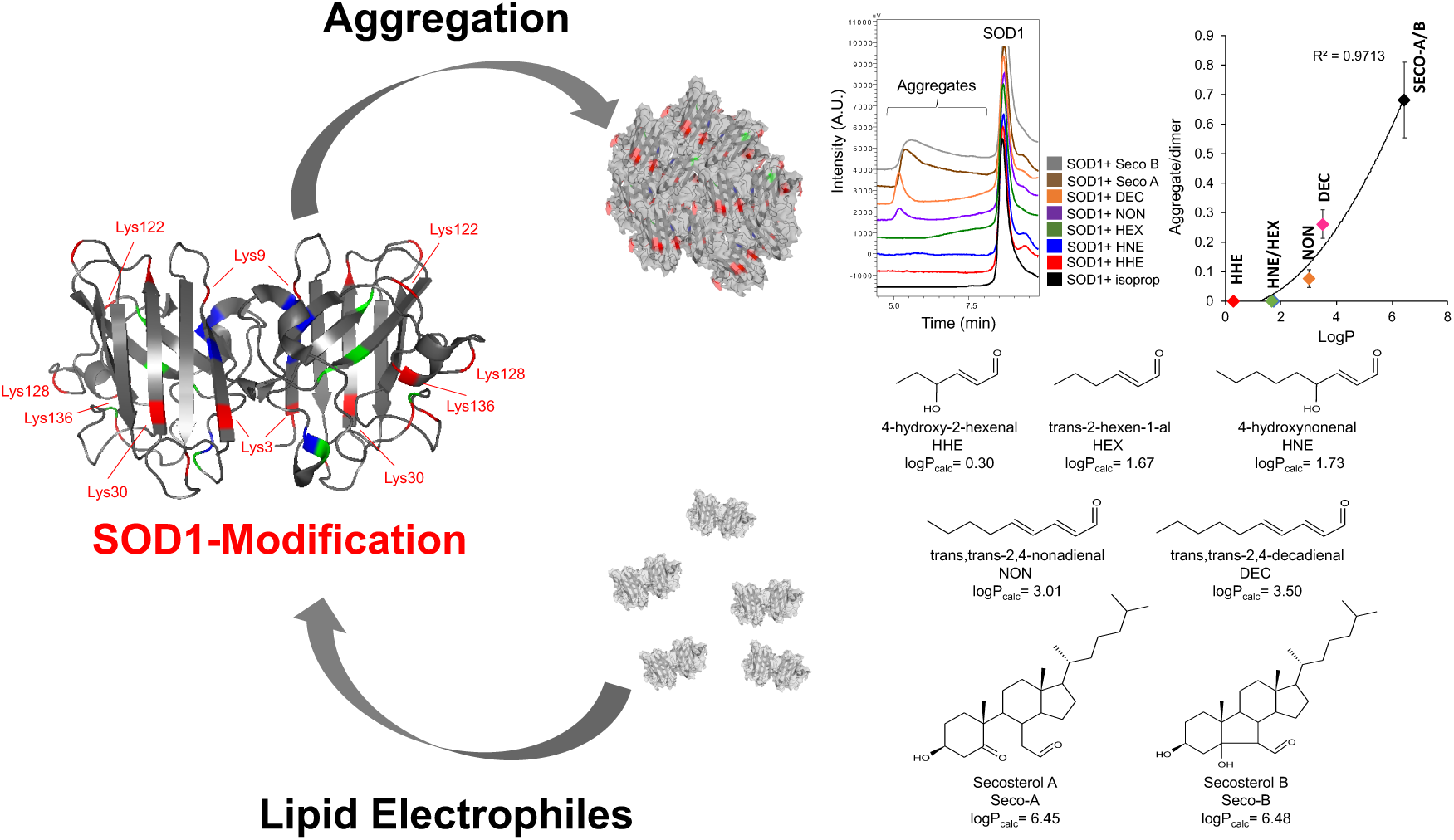

*Highlights:* - Aldehyde hydrophobicity is positively correlated to SOD1 aggregation; - Lys residues located nearby the SOD1 dimer interface and electrostatic loop are ubiquitously modified by all aldehydes; - Hydrophobic aldehydes increase the lipophilic potential surface of the region where they bind;

## INTRODUCTION

Oxidative stress has been regarded to play an important role in neurodegenerative diseases^1, 2^. Increased concentrations of redox biomarkers, including lipid peroxidation products and oxidized proteins have been detected in blood and brain of patients and rodent models of Alzheimer’s disease, Parkinson’s disease, Amyotrophic Lateral Sclerosis (ALS) and other neurological disorders.^3–8^ Lipids, which are usually highly unsaturated in neuronal cells, are susceptible to oxidation by enzymatic and non-enzymatic oxidation involving reactive oxygen species.^9–11^ In particular, lipid peroxidation leads to the formation of several electrophilic carbonyl compounds that are capable of modifying proteins and other components of the cell. ^12–14^

Lipid electrophiles are known to react with proteins and DNA, generally leading to irreversible damage through covalent adduction.^14–16^ Among lipid aldehydes, the better investigated are acrolein, malondialdehyde (MDA), 4-hydroxy-2-hexenal (HHE) and 4- hydroxy-2-nonenal (HNE).^17–19^ The α,β-unsaturated aldehydes can form adducts with nucleophilic residues of proteins by two different mechanisms: (1) Michael addition at the double bond to Lys, His or Cys residues or (2) Schiff base formation of the carbonyl group to Lys residues.^12, 14^ Lipid-derived aldehydes have already been shown to modify several proteins,^20, 21^ including those related to neurodegenerative diseases: α-synuclein,^22–25^ β-amyloid peptide^26, 27^ and superoxide dismutase (SOD1).^28^

Cholesterol is another lipid present in massive amounts in neurons and glial cells.^29^ Dysregulated cholesterol metabolism^30^ and massive accumulation of choleteryl esters have been reported in ALS^31^. Cholesterol undergoes oxidative damage, yielding hydroperoxides/hydroxides, ketones, epoxides as well as highly hydrophobic aldehydes named cholesterol secosterol aldehydes (Seco-A and Seco-B).^11, 32–35^ These hydrophobic aldehydes react mainly by Schiff base formation with Lys residues^36^ and their reactivity toward α-synuclein,^37^ β-amyloid peptide^38, 39^, myelin basic protein (MBP)^40^ and SOD1^41^ was previously investigated. Recently, secosterol aldehydes were detected in blood plasma and neural tissues from ALS rats.^41^

Mutations in the SOD1 gene have been linked to the development of familial ALS. The formation of high molecular weight SOD1 oligomers are implicated in both sporadic and familial ALS cases.^42–44^ Yet, the molecular mechanism leading to the formation of SOD1 aggregates is still not clearly understood. Some studies suggest that lipids, in particular, lipid peroxidation and its electrophilic products may play an important role in motor neuron degeneration^45^ and SOD1 aggregation.^46, 47^ Previously we reported that secosterol aldehydes are present in tissues isolated from ALS rat model and are effective in inducing SOD1 aggregation^41^. Here we aimed to compare the ability of lipid-derived electrophiles with different hydrophobicities to induce SOD1 modification and aggregation *in vitro*. SOD1 incubations with aldehydes showed that aggregation was positively correlated to their calculated hydrophobicities (LogP). Protein modification sites were mapped through LC-MS/MS analysis of the tryptic peptides. All aldehydes ubiquitously modified Lys residues located mainly at K3, the dimer interface (K9, K30) and the electrostatic loop (K122, K128 and K136). Interestingly, the less hydrophobic α,β-unsaturated aldehydes, HHE and HNE were likely more reactive, inducing modifications on Lys, His and Cys residues. However, they did not induce aggregation. Altogether, our study highlights the importance of electrophile hydrophobicity as a critical factor influencing protein aggregation.

## MATERIALS AND METHODS

### Chemicals

4-hydroxy-2-hexenal (HHE) and 4-hydroxy-2-nonenal (HNE) were purchased from Cayman Chemical (Ann Arbor, MI). Trans-2-hexen-1-al (HEX), *trans,trans*-2,4-nonadienal (NON) and *trans,trans*-2,4-decadienal (DEC) were purchased from Sigma (St. Louis, MO). 3β-hydroxy-5-oxo-5,6-secocholestan-6-al (Secosterol A, SECO-A) was synthetized by ozonization and purified as described by Wang and colleagues.^48^ 3β-hydroxy-5β-hydroxy-B-norcholestane-6β-carboxaldehyde (Secosterol B, SECO-B) was synthetized by photoxidation and purified as described by Uemi and colleagues.^33^ SOD1 (superoxide dismutase 1) was expressed in *Escherichia coli*, purified and its apo form prepared as described previously.^41^ Unless otherwise stated all chemicals were of the highest analytical grade and were purchased from Sigma, Merck or Fisher.

### Lipid aldehyde hydrophobicity determination

Aldehyde hydrophobicity was evaluated through its LogP (LogP_calc_) values calculated by the MoKa™ software,^49^ using the 3D structures generated by Corina v.3.20,^50^ and compared to those calculated by the Volsurf+ v.1.0.7.l software.^51^

### Incubations of SOD1 with lipid aldehydes

Apo form of SOD1 (10 μM) was incubated in 50 mM phosphate buffer, pH 7.4, containing 150 mM NaCl and 100 μM DTPA in the presence of 250 μM HHE, HNE, HEX, DEC, SECO-A or SECO-B at 37 °C during 24 h under gentle agitation. Isopropanol was the employed aldehyde solvent; therefore, 10% isopropanol was used as the control.

### SOD1 aggregate formation analysis

Ten μL of each incubation was analyzed by size-exclusion chromatography (SEC) using fluorescence detection with excitation wavelength at 280 nm and emission at 340 nm. Samples were eluted with 50 mM phosphate buffer, pH 7.4, containing 150 mM NaCl in the column BioSep-SEC-S3000 (300 × 7.8 mm, Phenomenex, USA) at 0.5 mL/min. Aggregate formation was evaluated and quantified by the ratio between the area of aggregates and the area of SOD1 dimer.

### Enzymatic digestion of SOD1

Incubations of SOD1 with aldehydes were first reduced with 5 mM sodium borohydride (NaBH_4_) for 1h at room temperature to stabilize the Schiff base adducts. Then samples were treated with 5 mM DTT (dithiotreitol) for 30 min at 60 °C to reduce disulfide bonds, followed by alkylation of Cys residues with 15 mM iodoacetamide for 30 min at room temperature. After that, SOD1 samples were digested with proteomic grade trypsin (Promega) for 18h in a 1:100 (w/w) ratio at 37 °C in the presence of RapiGest SF Surfactant (Waters).

### Characterization of modified peptides

Peptides resulting from tryptic digestion were analyzed using a nanoAcquity UPLC system (Waters, United States) with an ACQUITY UPLC-C18 column (20 mm × 180 μm; 5 μm) coupled to a TripleTOF 6600 mass spectrometer (Sciex, United States) as previously described.^41^ The Analyst TF® software (Sciex, United States) was used for data acquisition and the PeakView® software (Sciex, United States) for data processing. For analysis of protein sequence and modification, the Mascot® software (Matrix Science Ltd., London, United Kingdom) was used. The database in Mascot® search was SwissProt (accessed in 07/29/2018). The modifications were searched based on the possibilities of Schiff base formation to Lys residues after reduction with NaBH_4_ (HHE: +98.0731 Da; HNE: +140.1201 Da; HEX: +82.0782 Da; NON: +122.1095 Da; DEC: +136.1252 Da; and SECO-A/B: +402.3497 Da) as well as Michael addition to Lys, His and Cys residues (HHE: +114.0680 Da; HNE: +156.1150 Da; HEX: +98.0731 Da; NON: +138.1044 Da; DEC: +152.1201 Da; and SECO-A/B: +400.3341 Da). Carbamidomethyl (+57.0214 Da) was also considered as a possible modification of Cys residues. The mass tolerance was 10 ppm for MS experiments and 0.05 Da for MS/MS experiments. Up to 4 trypsin missed cleavages were considered in the search. Modified peptides identified by Mascot® were further validated by manual identification of the sequence (Supplementary MS/MS spectra are provided).

### Covalent docking procedures

#### Ligand structures preparation for docking purpose

Prior to docking runs, three-dimensional ligand structures for, respectively, HHE, HEX, HNE, NON and DEC were generated using Corina v.3.20.^50^ As compounds HNE and HHE were used in the experimental assays as racemic mixtures, both enantiomeric forms were generated and used for docking. Seco-A and seco-B 3D structures were obtained from *PubChem Bioassay Tools* (available online on https://pubchem.ncbi.nlm.nih.gov/). The 3D structure of each aldehyde was minimized using the Sybyl-X v.2.1.1 software (Certara L.P., St Louis, MO). Energy minimization was done according to the Powell method, being stopped when the energy difference between interactions was lower than 0.05 kcal.mol^-1^.Å-1.

#### Protein structure preparation for docking purpose

For the same purpose, protein structure (SOD1, PDB: 3ECU, 1.9 Å resolution) was also prepared using Sybyl-X v.2.1.1. All hydrogen atoms were added, and predominant protonation states for amino acids side chains were set at pH 7.4. All water molecules were removed from the original pdb file.

#### Covalent docking procedure

Covalent docking was performed using GOLD docking suite v.5.2 (The Cambridge Crystallographyc Data Centre).^52^ GOLD program assumes that there is just one atom linking the ligand to the protein. An angle-bending potential has been incorporated into the fitness function to ensure that the geometry of the bound ligand is correct.^53, 54^ For each docking run, zeta nitrogen atom of Lys side chain (NZ) was defined as the link atom for the covalent bond. Each binding site was defined as all residues with at least one heavy atom within 10 Å from the NZ atom of the corresponding Lys residue. Ten docking runs were performed for each ligand, using default settings and scoring function ChemPLP.^55^

### Lipophilic potential surfaces generation

Lipophilic potential surfaces for SOD-1 were generated using MOLCAD^56^ module of Sybyl-X v.2.1.1 program (Certara L.P., St Louis, MO), applying the Connolly method.^57^ All generated surfaces were qualitatively analyzed by visual inspection.

## RESULTS

### Aldehyde-induced SOD1 aggregation increases exponentially with their LogP values

To compare the ability of lipid-derived electrophiles to promote SOD1 aggregation, we selected seven biologically relevant aldehydes: 4-hydroxy-2-hexenal (HHE) and 4-hydroxy-2-nonenal (HNE), trans-2-hexen-1-al (HEX), *trans,trans*-2,4-nonadienal (NON), *trans,trans*-2,4-decadienal (DEC), 3β-hydroxy-5-oxo-5,6-secocholestan-6-al (Seco-A) and 3β-hydroxy-5β-hydroxy-B-norcholestane-6β-carboxaldehyde (Seco-B) (Figure 1). The LogP of these aldehydes was estimated using MoKa™ software.^49^ Theoretically estimated LogP values ranged from 0.30 (HHE) to ∼6.5 (Seco-A and Seco-B). Aldehyde stock solutions were prepared in isopropanol and incubated with metal-free SOD1 (apo-SOD1) as previously described.^41^ We used apo-SOD1 because several lines of evidence indicate that this immature form plays a key role in aberrant oligomerization^58, 59^. After 24 h, high molecular weight protein aggregates were analyzed by size-exclusion chromatography (SEC). In this analysis, SOD1 dimers appeared as a single peak at 8.5 min and large SOD1 aggregates appeared as a broad peak between 5 to 7.5 min in incubations containing NON, DEC and Seco-A or Seco-B (Figure 2A). To quantify the amount of protein aggregates, the peak areas corresponding to the dimer and aggregate were integrated and expressed as aggregate/dimer area ratio. This area ratio was plotted against estimated aldehyde hydrophobicity (LogP, Figure 2B). Interestingly, this plot showed a strong positive correlation between aldehyde hydrophobicity and protein aggregation. Under our experimental conditions, the less hydrophobic lipid electrophiles (LogP<2), HHE, HNE and HEX, did not induce detectable SOD1 aggregation. In contrast, SOD1 aggregation was dramatically increased by the more hydrophobic electrophiles (LogP>3) in the following order: nonadienal (NON) < decadienal (DEC) < secosterol B (SECO-B) (Figure 2B).

**Figure 1.**
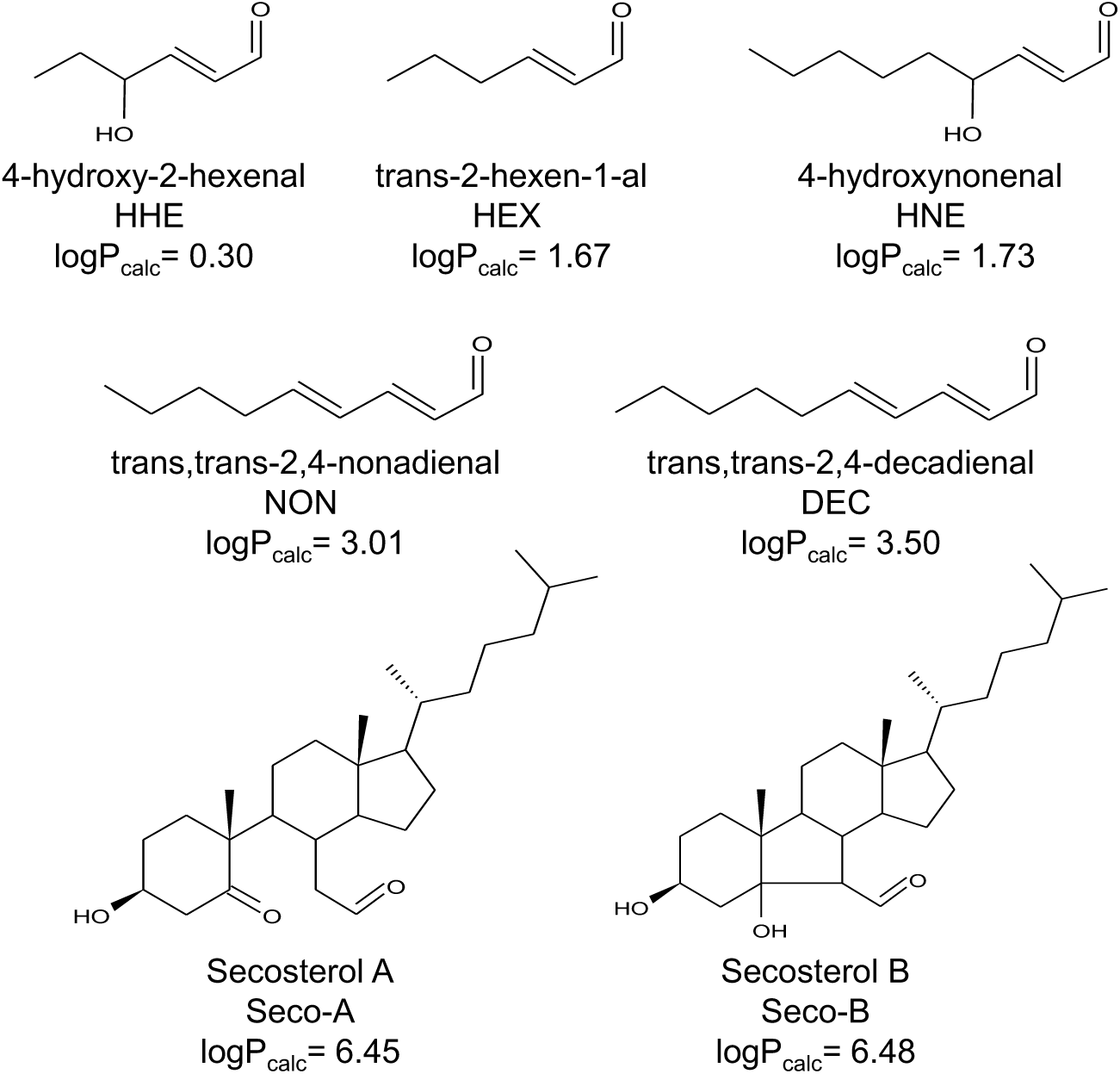
Structure of lipid-derived electrophiles and their corresponding partition coefficients (logP_calc_) calculated by the Moka™ software.

**Figure 2.**
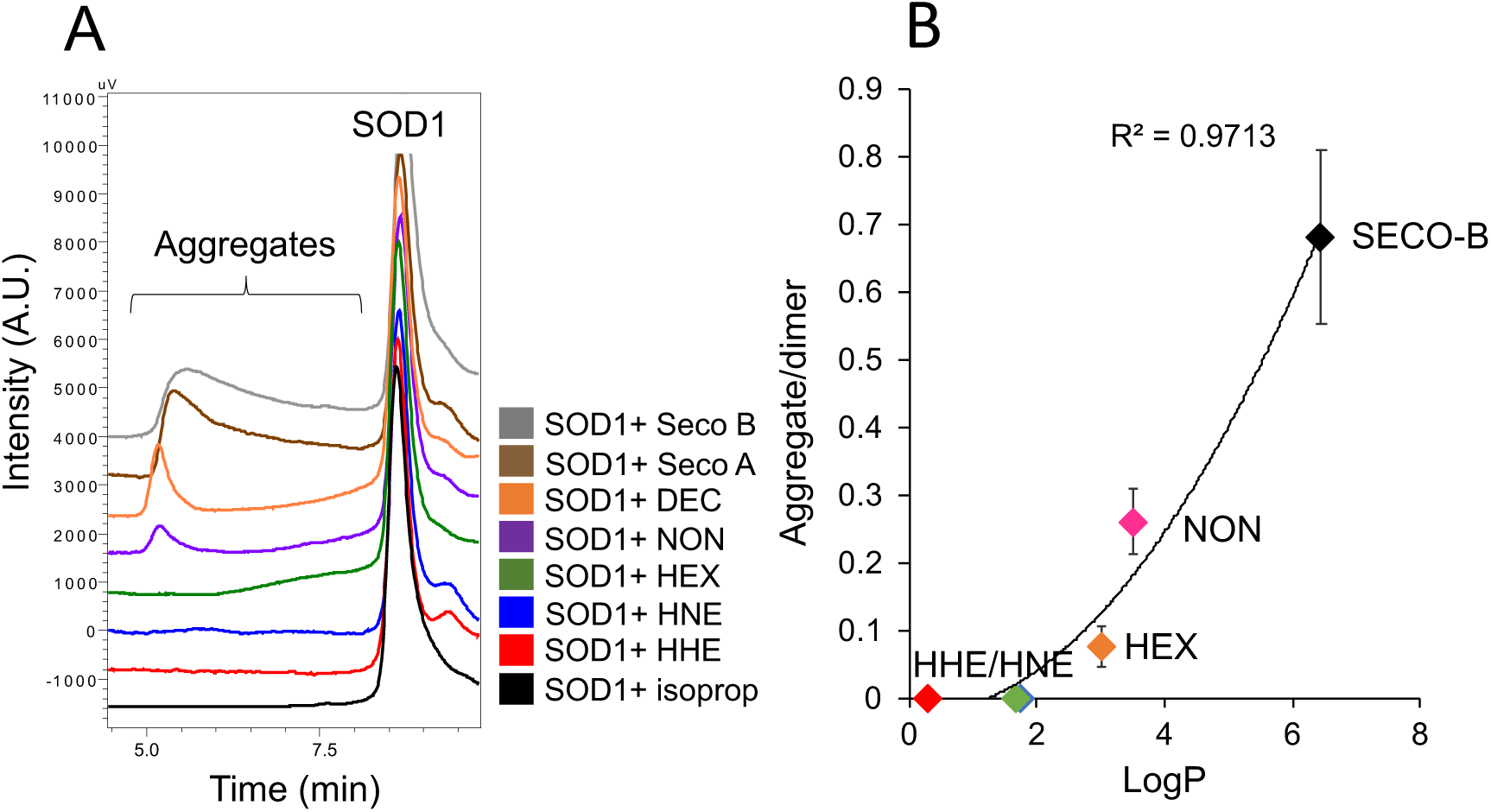
Correlation between SOD1 aggregation and lipid electrophile hydrophobicity (logP). (A) Size-exclusion chromatography (SEC) analysis of SOD1 after incubation with different lipid electrophiles. 10 μM apo-SOD1 was incubated with 250 μM aldehydes at 37°C for 24h. Aliquots of 10 μL of each incubation were analyzed by SEC using fluorescence detection with excitation wavelength at 280 nm and emission at 340 nm. (B) Correlation between SOD1 aggregate formation (aggregate area/dimer area) and lipid aldehyde hydrophobicity (calculated logP values). Correlation showed a R^2^ = 0.977.

### Characterization of aldehyde adduction sites in SOD1

To characterize the residues modified by lipid electrophiles, proteins incubated with HHE, HNE, HEX, NON, DEC or SECO-A/B were subjected to trypsin digestion followed by peptide sequencing analysis on a reverse-phase nLC-ESI-MS/MS. The obtained MS/MS data were sequenced by Mascot® using the bottom-up approach. Covalently modified peptides were searched according to two possibilities, Schiff-base or Michael-type adducts. Each modified peptide sequence was confirmed manually by the analysis of the MS/MS spectra (Supplementary MS/MS spectra). Protein electrophile adduction with secosterol aldehydes (SECO-A/B) occurred exclusively by Schiff-base formation, while both Schiff-base and Michael adducts were observed for the α,β-unsaturated aldehydes (Table 1).

**Table 1.**
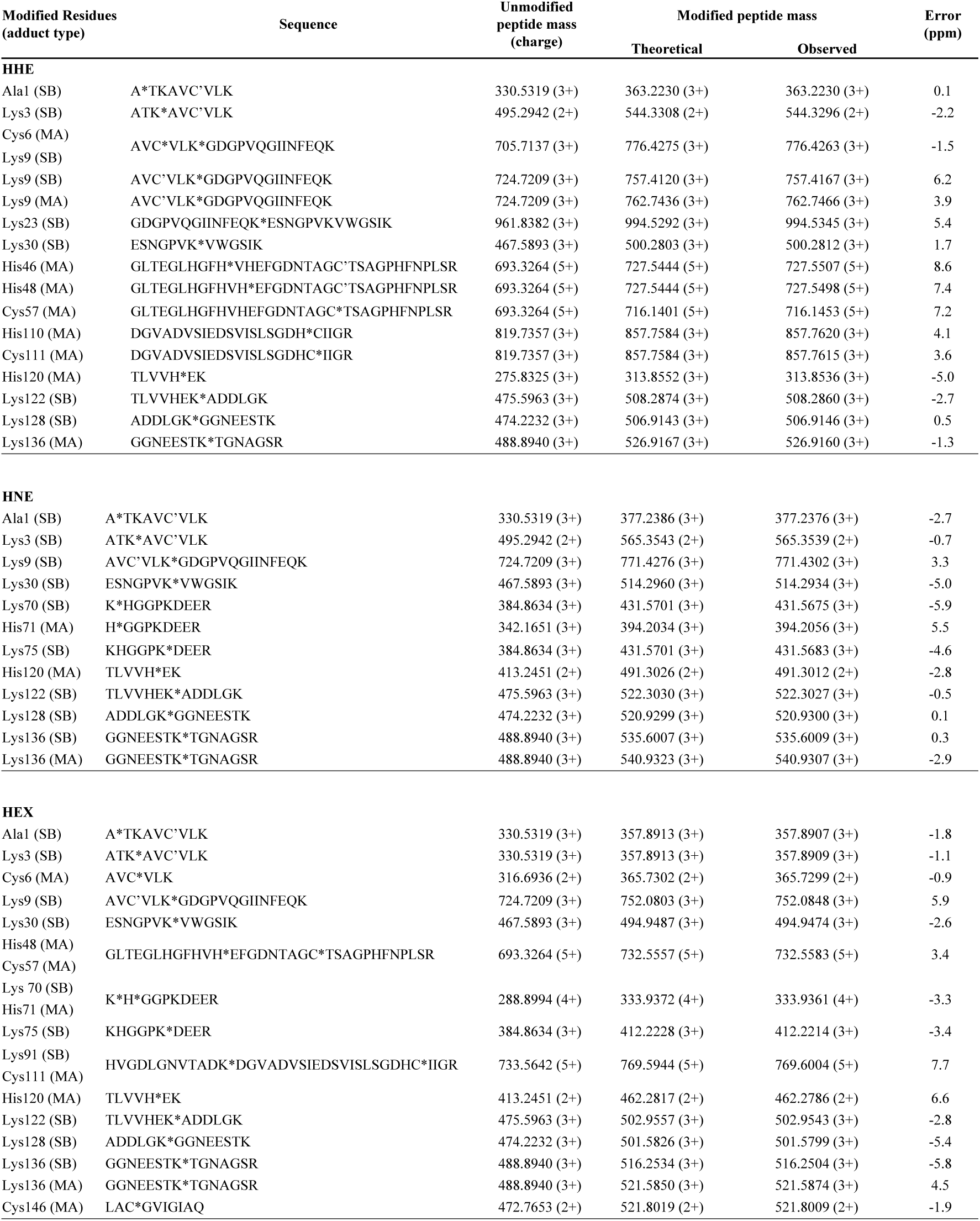
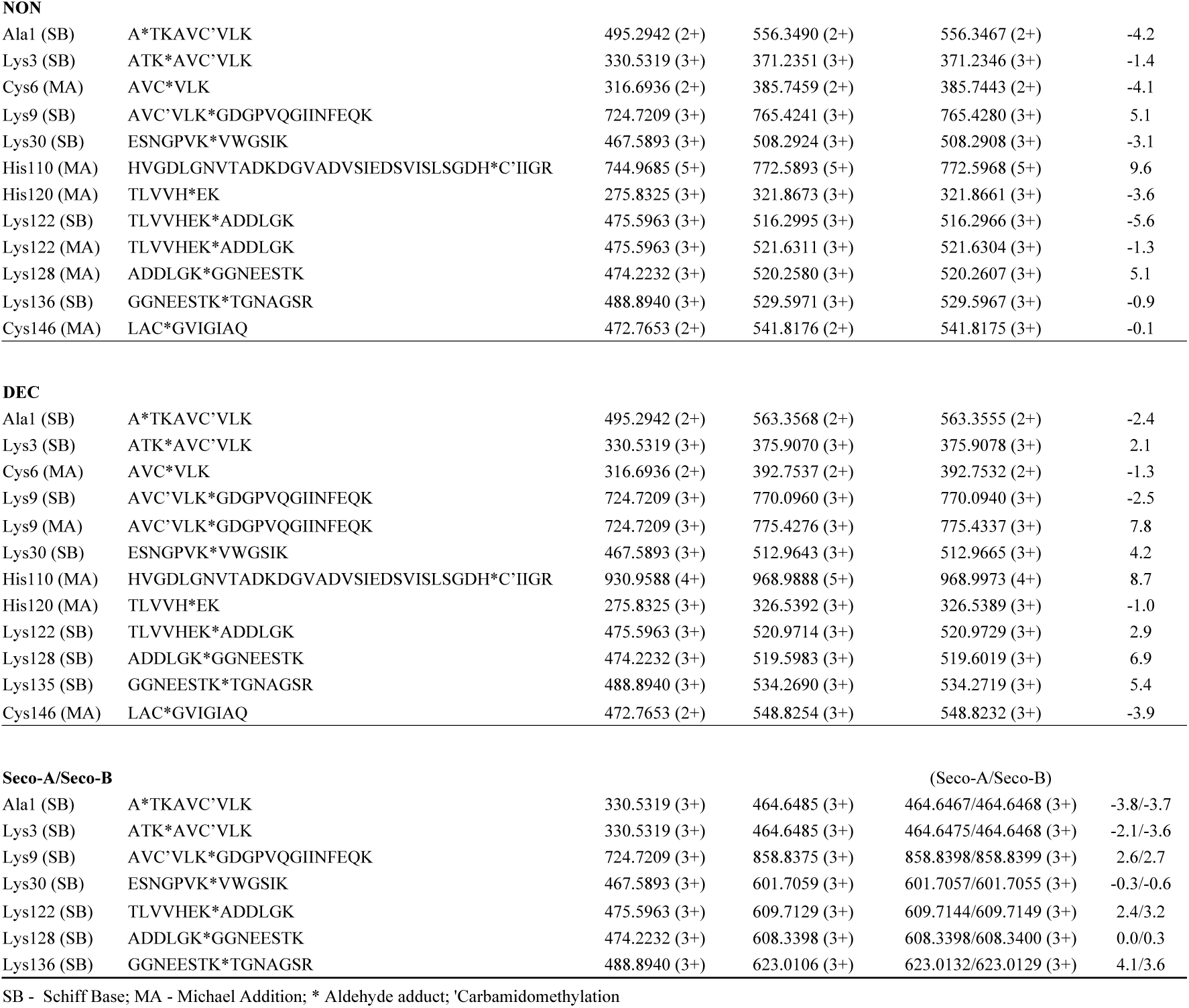
SOD1 modified residues identified after incubation with HHE, HNE, HEX, NON, DEC or SECO-A/B.

The position of the modified residues in SOD1 structure is summarized in Figure 3. Interestingly, from the 11 Lys present in SOD1, only 6 were commonly modified by all tested aldehydes: K3, K9, K30, K122, K128 and K136 (Figure 3A). Modified Lys 3 and 9 residues are located nearby the dimer interface and Lys 122, Lys 128 and Lys 136 are residues located at Loop 7 or electrostatic loop (Figure 3B). In contrast, the less solvent-exposed Lys residues were selectively modified only by the less hydrophobic aldehydes (HHE, HNE and HEX, Log P<2). Lys 23 was modified only by HHE, and Lys 70 and 75 were modified by HNE and HEX. Interestingly, similarly to the results obtained for the aggregation assay, the retention time of the adducted peptides during reversed-phase nLC-MS/MS analysis was positively correlated to aldehyde hydrophobicity (Figure 4 and Figure 1S). Peptides modified by the less hydrophobic aldehyde, HHE, eluted earlier at around 20 min, while those modified by the more hydrophobic aldehydes had their retention time increased, being retained up to 60 min with Seco-A/B.

**Figure 3.**
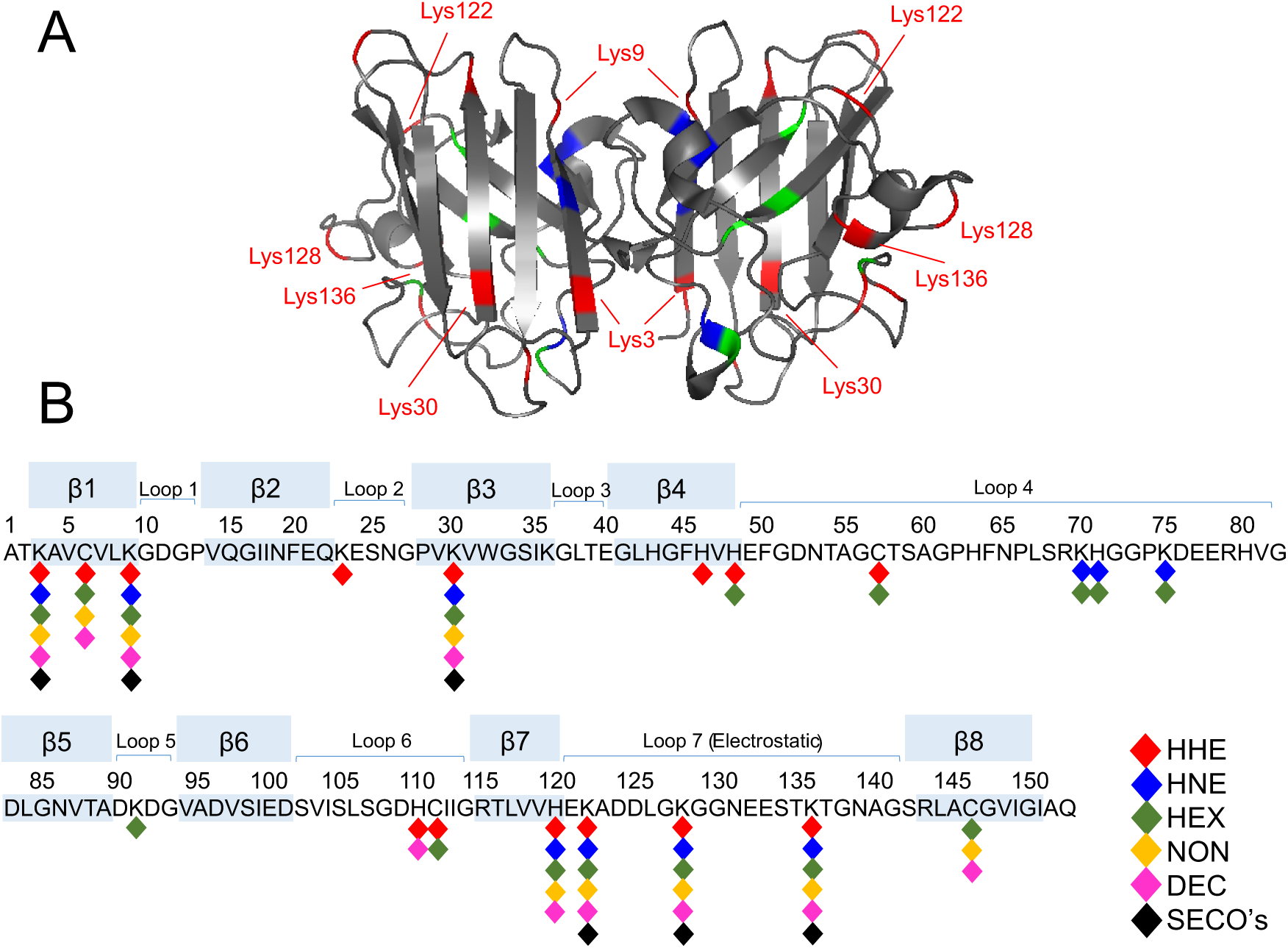
SOD1 residues modified by the lipid electrophiles. (A) Protein quaternary structure of SOD1 (PDB code 3ECU). Lys residues are highlighted in red, cysteine residues in blue and histidine residues in green (B) Linear sequence of SOD1. Modified residues are attached to the representative color for each aldehyde (colored diamonds).

**Figure 4.**
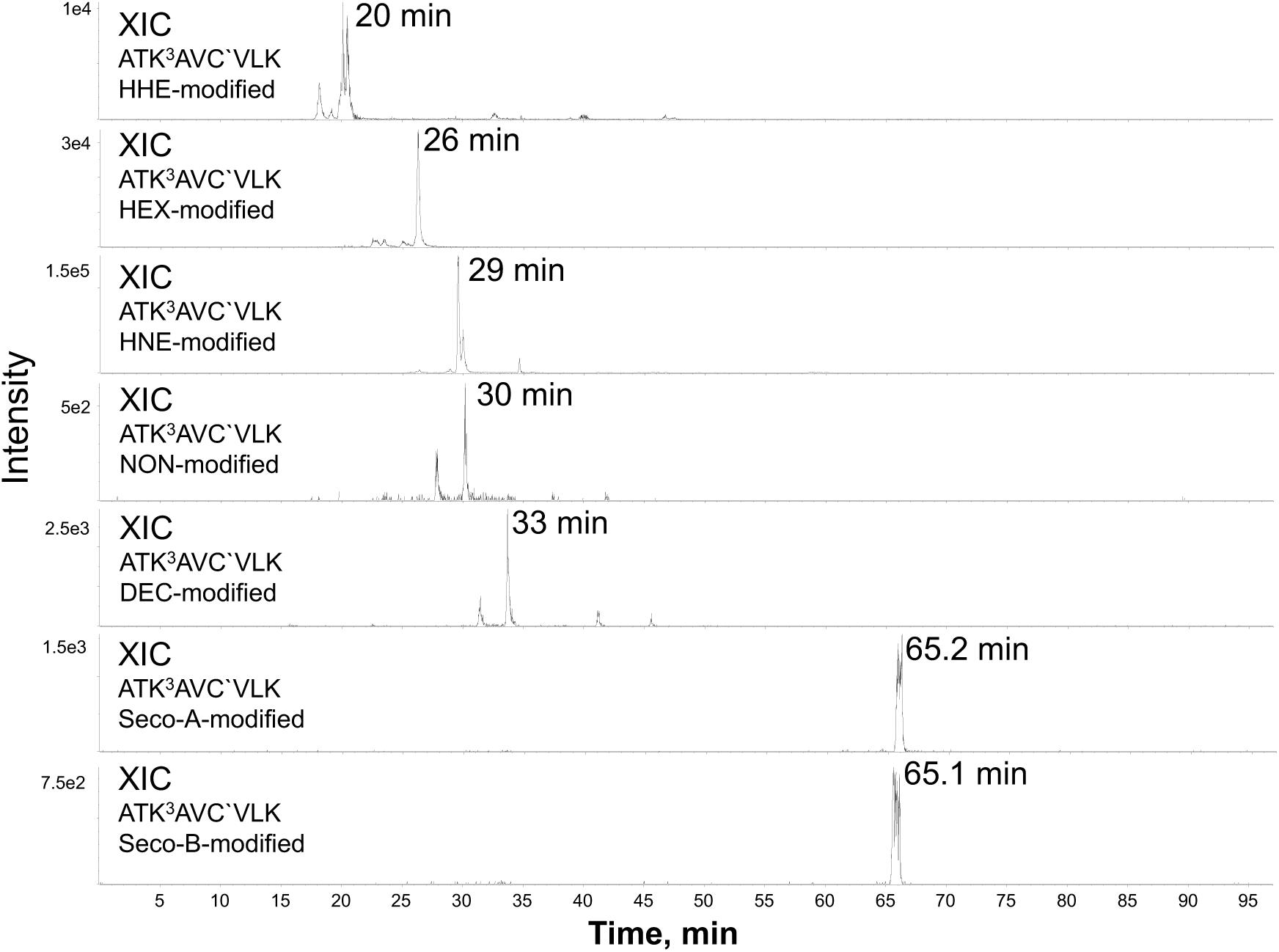
Increased retention time of modified peptides. Peptides resulted from tryptic digestion were analyzed using a nanoAcquity UPLC system with an ACQUITY UPLC-C18 column (20 mm × 180 μm; 5 μm) coupled to a TripleTOF 6600 mass spectrometer (Sciex, United States). Figure represents the extracted ion chromatogram (XIC) for the peptide ATK^3^AVCVLK modified by each aldehyde.

In addition to Lys residues, His and Cys residues were modified by all the α,β-unsaturated aldehydes (Figure 3B). Among His residues, H120 was found modified by all the α,β-unsaturated aldehydes (HHE, HNE, HEX, NON, DEC), whereas other His residues (H46, H48, H71, H75, H110) were selectively modified only by smaller and less hydrophobic aldehydes (HHE, HNE and HEX). Among Cys residues, C6 was modified by most α,β-unsaturated aldehydes, except by HNE. C111 and C57 were modified only by HHE and HEX, whereas C146 was modified only by the 2,4-alkadienals, HEX, NON and DEC (Figure 3B).

Next, to estimate the extension of protein modifications induced by the aldehydes we compared the relative percentage of intact peptides detected in the control and in aldehyde-treated samples (Figure 5). HNE and HHE showed to be the two most reactive aldehydes, leading to the consumption of more than 50 % of the peptides. HEX, NON and DEC presented an intermediate reactivity, leading to 20-50 % of consumption of the peptides. The secosterol aldehydes (Seco-A and Seco-B) were the less reactive, consuming less than 10 % of the peptides in most cases (Figure 5). This comparison indicates that, although Seco-A/B were likely to induce fewer modifications, they were the most effective ones to promote protein aggregation (Figure 2). Conversely, the less hydrophobic and reactive short-chain aldehydes, HHE and HNE, induced more modifications but were less efficient in inducing SOD1 aggregation. Therefore, altogether this data strongly indicates that aldehydes’ hydrophobicity determines their efficiency in inducing protein aggregation, at least under *in vitro* conditions.

**Figure 5.**
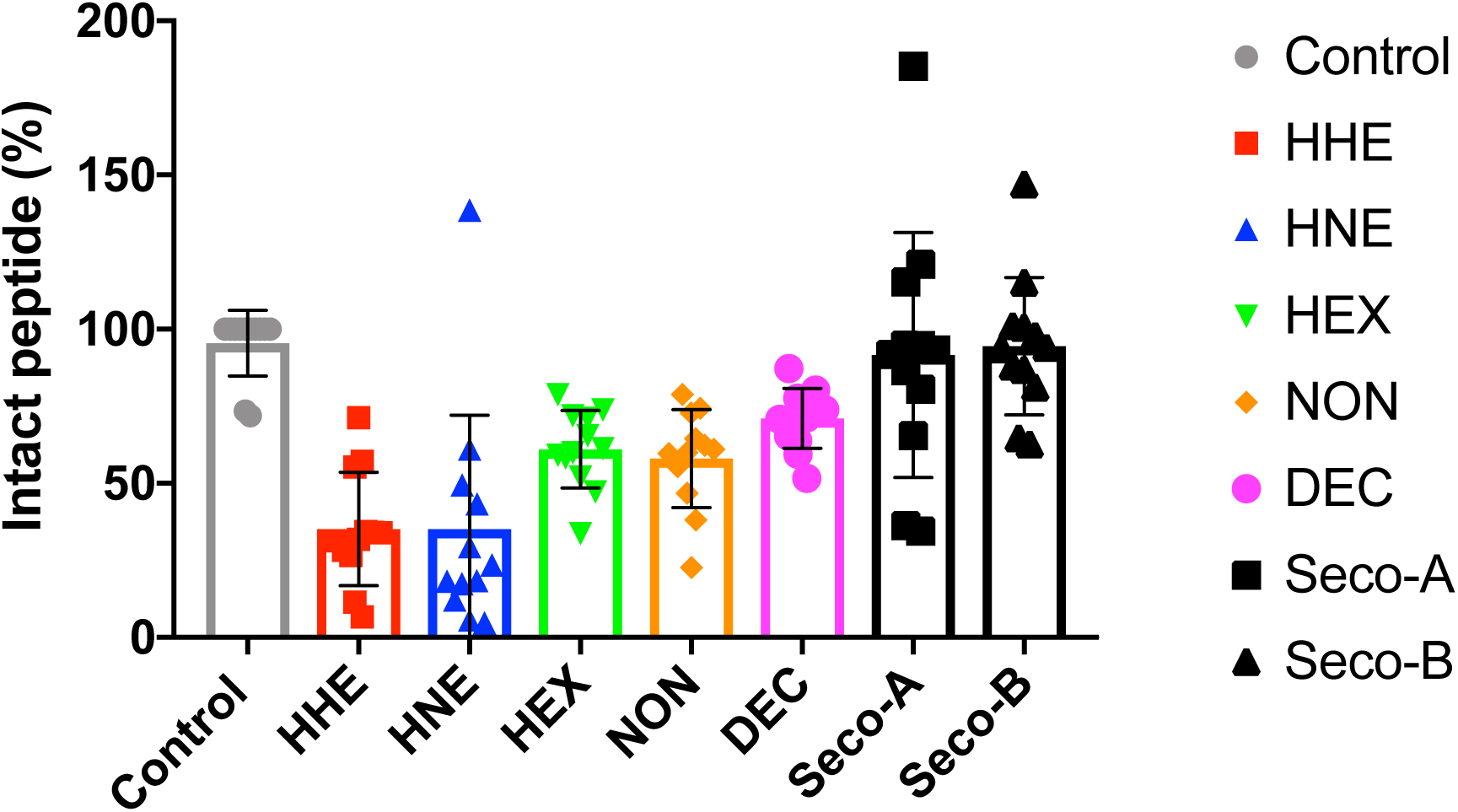
Estimated amount of intact SOD1 peptides found after incubation with lipid-derived electrophiles. Areas of the peptides resulted from tryptic digestion were analyzed using MultiQuant® software. Data represent the sum of areas of all unmodified peptides normalized by the area of total ion chromatogram (TIC). Results were converted in percentage relative to the control.

### Docking poses and lipophilic surface calculations

In order to get structural insights into the protein-ligand interactions that seem to occur between the aldehyde molecules and the residues surrounding Lys3, Lys9, Lys30, Lys122, Lys128 and Lys136 (main residues modified by the aldehydes), covalent dockings were performed, using GOLD. ChemPLP scores for the best ranked poses for each docked aldehyde are shown in Table S1. In general, the higher scores, indicating the highest number of both hydrophobic contacts and hydrogen-bond interactions, were observed for the secosterol aldehydes (Seco-A/B). On the other hand, the lowest scores, indicating fewer hydrophobic and hydrogen bond interactions, were found with the less hydrophobic aldehydes, HHE and HNE. This scoring algorithm suggests that protein-aldehyde interactions involving the specified Lys residues are favored for the more hydrophobic aldehydes.

Ligand-protein interaction diagrams, showing the main non-covalent interactions identified between the docked aldehyde molecules and some amino acids surrounding each lysine residue, were also generated (Table S2). Most of the identified interactions corresponded to hydrophobic contacts established with some nonpolar residues, such as Val5, Ile17 and Ala152 near to Lys3; Ile17, Val7 and Val94 near to Lys9; Trp32 near to Lys30; Ala123 and Leu42 near to Lys122; Leu126 near to Lys128 and Thr137 near to Lys136.

Considering that Seco-A and Seco-B were the most effective aldehydes in inducing SOD1 aggregation, the best ranked predicted docking poses of these aldehydes linked to Lys residues (Lys3, Lys9, Lys30, Lys122, Lys128 and Lys136) were further analyzed. Lipophilic potential surfaces were calculated using MOLCAD, in which the surface colors range from brown (highest lipophilic area of surface) to blue (highest hydrophilic area of surface) (Figure S2). Interestingly, a contrasting effect on the lipophilic surface was apparent when Seco-A/B are covalently attached to a predominantly hydrophilic region, such as the Lys136 surrounding area (see Figure 6). As can be noticed, Seco-A adduction to Lys 136 led to increased SOD1 lipophilic surface (brown colored surface, Figure 6H) compared to HHE (Figure 6C), in agreement with its higher capacity to induce protein aggregation.

**Figure 6.**
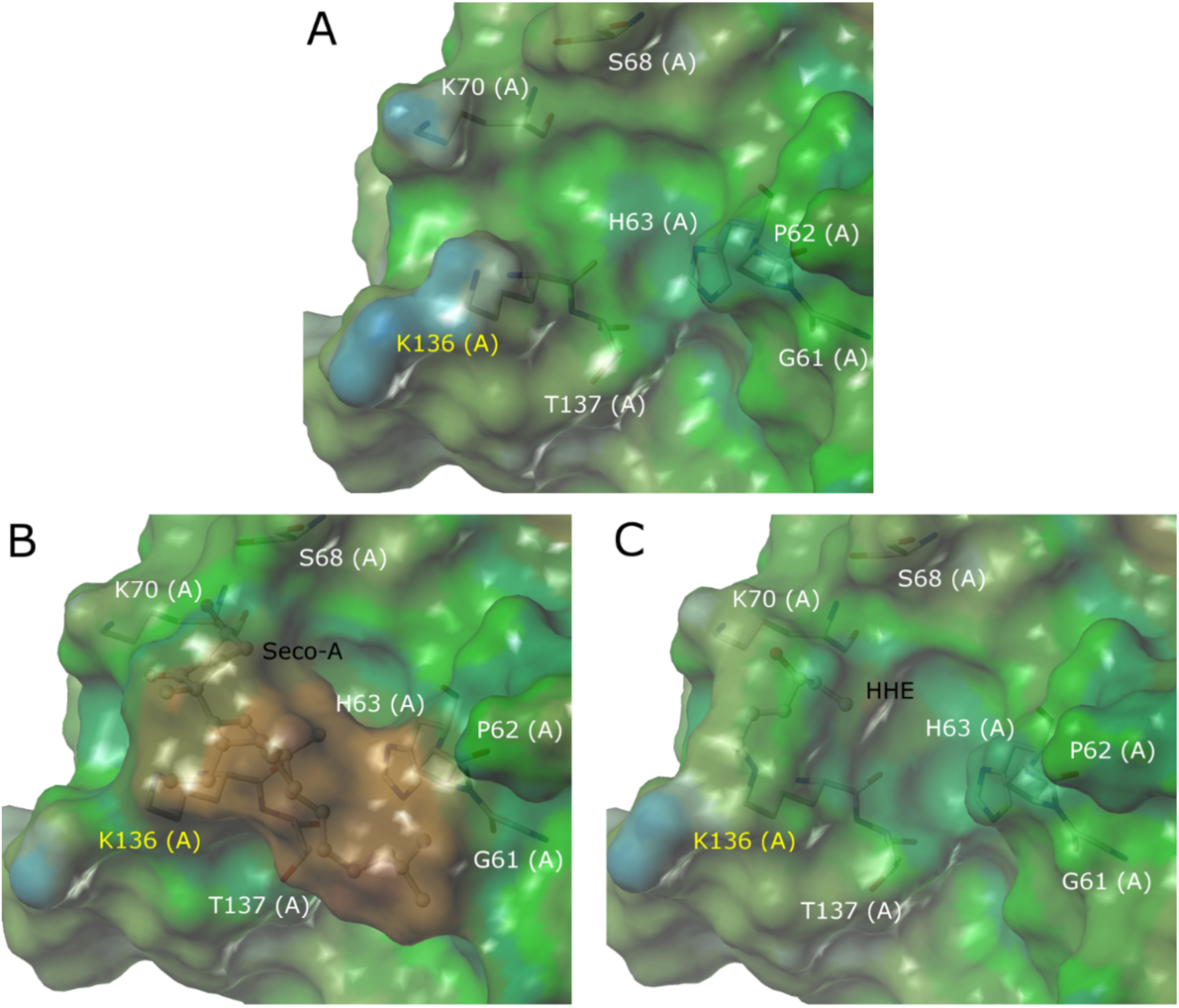
Lipophilic potential surfaces for SOD1 calculated using MOLCAD. (A) Lipophilic potential surface for the binding site around Lys136; (B) Lipophilic potential surface for seco-A covalently bound to Lys136; (C) Lipophilic potential surface for HHE covalently bound to Lys136. The color ramp for lipophilic potential ranges from brown (highest lipophilic area of surface) to blue (highest hydrophilic area of surface). Seco-A and HHE are shown as balls and sticks, and amino acid residues are shown as sticks. Hydrogen atoms are omitted for clarity.

## DISCUSSION

Protein misfolding and aggregation in the central nervous system are associated with neurodegenerative disorders^60^. Much attention has been focused on the role of reactive electrophilic lipids or reactive carbonyl species (e.g. HNE) generated under conditions of elevated oxidative stress^4, 8, 24^. These electrophilic compounds exhibit high reactivity toward nucleophilic groups in proteins and other macromolecules, leading to their covalent modification. Advanced mass spectrometry profiling methods have expanded our knowledge on the pathological and physiological impact of lipid electrophiles on different cellular targets ^20, 61–63^. A great diversity of protein adducts and cross-links have been characterized and collectively termed as advanced lipoxidation end products.^64, 65^ Here, we compared the ability of lipid electrophiles with different hydrophobicities to promote SOD1 aggregation. Interestingly, our data shows that aldehyde-induced SOD1 aggregation is highly correlated to their hydrophobicity.

Previous studies have investigated SOD1 modifications by electrophiles, such as acrolein and advanced glycation end products. In the study by Kang,^28^ high concentrations of acrolein (mM) showed to inhibit SOD1 activity by modification of Ser, His, Arg, Thr and Lys residues. Kato and co-workers^66^ detected SOD1 aggregates imunoreactive to advanced glycation end-product’s modification site in ALS patients and mice expressing human SOD1-G85R mutation. Both studies suggested that electrophiles may play significant roles in SOD1 homeostasis. Here, we not only demonstrated that lipid-derived electrophiles can modify and induce SOD1 aggregation, but also that the effect is dependent on the electrophile hydrophobicity (Figure 2). The data corroborate our view that SOD1 modification by aldehydes increases the hydrophobic surface of the protein, increasing their aggregation propensity. Likewise, Liu and co-workers^27^ found out that β-amyloid peptide misfolding and fibrillogenesis are promoted by HNE but not by HHE, which is a less hydrophobic aldehyde.

It is important to notice that SOD1 aggregation efficiency did not correlate to the reactivity of the aldehyde, but more critically to its hydrophobicity. Indeed, HHE and HNE induced a higher degree of protein modification compared to other aldehydes (Figure 5), but did not induce their aggregation (Figure 2). On the other hand, NON, DEC, and SECO-A/B induced fewer modification, but were the most effective inducers of SOD1 aggregation, a property that showed to be correlated to their higher hydrophobicity. Analysis of the modified residues revealed that all aldehydes induced modification on specific Lys residues located primarly at K3 and K9, nearby the dimer interface, and at K122, K128 and K136, located within the electrostatic loop (residues 121-143). These two regions (dimer interface - loop IV and the electrostatic loop - loop VII) are frequently affected by genetic and non-genetic factors, the later including an array of post-translational modifications (PTM, e.g. acylation, sumoylation, ubiquitination, glycation)^67, 68^. Dimer interface modifications have been shown to enhance monomer formation^69^, increasing the production of high molecular weight inclusions^70^. Furthermore, disorders affecting the eletrostatic loop have been correlated with structural alterations leading to aggregation^71, 72^. Recently, a study by Mojumdar *et al.* provided additional evidences indicating that the region comprising electrostatic loop is the least stable part of the protein^70^, consistent with the notion that this region acts as a primary locus for misfolding. Thus, our study highlights the potential impact that lipid electrophile-induced Lys modifications occuring at these segments would have on SOD1 structural stability.

Lysine is a ubiquitous post-traslational modification site in the human proteome^73^. Acylation and acetylation of lysine residues in SOD1 have been previously described as potencial mechanisms to decrease the rate of protein nucleation and prion-like SOD1 aggregation.^74, 75^ These studies attributed the inhibitory effect of covalent modification to the increase of net negative surface potential and repulsion between SOD1 species. ^74^ Our data revealed that modifications promoted by the less hydrophobic aldehydes, HHE and HNE do not lead to protein aggregation. In analogy to the acetylation study, it can be hypothesized that Lys conjugation to these aldehydes could elicit a protective effect by blocking Lys charges. In this situation, small and less hydrophobic aldehydes (LogP<2) lead to a net increase in surface negative charge that blocks SOD1 aggregation. On the other hand, the size and polarity of the protein modifier seem to have a greater impact on the final outcome of protein aggregation. In this context, when the protein is modified by highly hydrophobic electrophiles, like the secosterol aldehydes (LogP ∼6), the net increase in protein hydrophobic surface greatly enhances their aggregation propensity.

Covalent docking analysis expanded our notion on how the aldehydes, in particular secosterol aldehydes, might be positioned/directed to specific Lys residues in SOD1 – the Lys3, Lys9, Lys30, Lys122, Lys128, and Lys136. Although there are other residues that were randomly modified by lipid electrophiles, including other lysines (Lys23, Lys70 and Lys75), histidines (His46, His48, His71, His110 and His 120) and cysteines (Cys6, Cys57, Cys111 and Cys146), the previously mentioned Lys residues were ubiquitously modified by all aldehydes, including the most hydrophobic and less hydrophobic ones. Detailed inspection of those Lys suroundings reveals that some hydrophobic residues, such as Val, Ala, Trp and Ile (Table S2), may play important roles directing aldehydes to the conjugation site. Hydrophobic residues may help recruit aldehydes to these areas by interacting with their hydrophobic surface. The enhancement in the lipophilic surface observed by binding of hydrophobic secosterol aldehydes to Lys136, a residue located at a relatively polar region (Figure 6), illustrates how ligand hydrophobicity affects protein surface hydrophobicity and increases protein aggregation.

In summary, there is an increasing number of studies arguing for the importance of oxidized lipids in neurodegenerative diseases^6, 7, 76^. In this context, our *in vitro* study shed light on a critical link between lipid peroxidation and protein aggregation, which is putatively associated with the pathology of these disorders. Our data specifically bear evidence that lipid electrophile hydrophobicity is critical to ligand-induced SOD1 aggregation. Given the massive abundance of cholesterol in brain tissues^29^ and the solid link between cholesterol metabolism and neurodegenerative diseases,^77, 78^ *in vivo* experiments with cells or animal models may provide additional clues on the role of highly hydrophobic secosterol aldehydes in protein aggregation.

## Acknowledgments

We would like to thank Fernando R. Coelho for technical assistance with SOD1 protein expression and purification. This work was supported by Fundação de Amparo à Pesquisa do Estado de São Paulo (FAPESP, CEPID-Redoxoma 13/07937-8 and 10/50891-0), Conselho Nacional de Desenvolvimento Cientifíco e Tecnológico (CNPq, Universal 424094/2016-9), NAP-Redoxoma, Pró-Reitoria de Pesquisa USP and CAPES. The Ph.D. scholarship of L.S.D. was supported by CNPq. The Ph.D. scholarships of L.G.V., A.I. and E.P. were supported by FAPESP (2014/07248-0, 2017/13804-1, 2012/06633-2). E.P. has a post-doctoral CAPES fellowship (88887.185840/2018-00).

